# Reciprocal lateral hypothalamic and raphé GABAergic projections promote wakefulness

**DOI:** 10.1101/2020.11.04.367722

**Authors:** Mary Gazea, Szabina Furdan, Péter Sere, Lukas Oesch, Benedek Molnár, Giuseppe Di Giovanni, Lief E. Fenno, Charu Ramakrishnan, Joanna Mattis, Karl Deisseroth, Susan M. Dymecki, Antoine R. Adamantidis, Magor L. Lőrincz

## Abstract

The lateral hypothalamus (LH), together with multiple neuromodulatory systems of the brain, such as the dorsal raphé nucleus (DR), is implicated in arousal, yet interactions between these systems are just beginning to be explored. Using a combination of viral tracing, circuit mapping, electrophysiological recordings from identified neurons and combinatorial optogenetics in mice, we show that GABAergic neurons in the LH selectively inhibit GABAergic neurons in the DR resulting in increased firing of a substantial fraction of its neurons that ultimately promotes arousal. These DR_GABA_ neurons are wake active and project to multiple brain areas involved in the control of arousal including the LH, where their specific activation potently influences local network activity leading to arousal from sleep. Our results show how mutual inhibitory projections between the LH and the DR promote wakefulness and suggest a complex arousal control by intimate interactions between long-range connections and local circuit dynamics.

## Introduction

Brain-wide neuronal activities are strongly modulated across sleep-wake states (Saper et al., 2010; Weber and Dan, 2016) attributable to the influence of neuromodulatory systems of the brainstem (Jones, 2020), basal forebrain (Fuller et al., 2011; Xu et al., 2015) and lateral hypothalamus (LH) (Bonnavion et al., 2016; Arrigoni et al., 2019). In this context, the LH stands both as an anatomical and a sleep-wake hub.

Various LH neuronal populations are known to modulate sleep-wake states (Saper et al., 2010; Weber and Dan, 2016), energy intake and reward (Bernardis and Bellinger, 1996; Stuber and Wise, 2016). Hypocretins/orexins (Hcrt/Ox) and Vgat-expressing neurons of the LH (LH_Vgat_) control wakefulness (Adamantidis et al., 2007; Carter et al., 2009; Carter et al., 2012; Herrera et al., 2016; Venner et al., 2016). Optogenetic activation of LH_Vgat_ neurons induced rapid arousal through direct modulation of the reticular thalamus and brainstem circuits in the locus coeruleus area (Herrera et al., 2016), concomittent to an inhibition of anterior hypothalamic sleep-promoting circuits (Venner et al., 2019). In contrast, activation of MCH neurons promotes REM, and to a lesser extent NREM, sleep (Verret et al., 2003; Jego et al., 2013; Konadhode et al., 2013; Tsunematsu et al., 2014). Yet, whether and how LH_Vgat_ neurons also modulate the activity of other brainstem neuromodulatory circuits remains unclear.

LH_Vgat_ neurons are reciprocally connected to multiple brainstem nuclei (Bonnavion et al., 2016), including the DR (Weissbourd et al., 2014). The DR is an appealing candidate for influencing sleep-wake states since its major neuromodulator serotonin rapidly influences sensory (Kapoor et al., 2016; Lottem et al., 2016), motor (Liu et al., 2014; Correia et al., 2017; Seo et al., 2019) and cognitive (Miyazaki et al., 2011; Liu et al., 2014; Cohen et al., 2015; Fonseca et al., 2015) functions. Importantly, the DR has been involved in the regulation of various patho-physiological functions including depression, which is often accompanied by sleep disturbances (Lowry et al., 2008; Steiger and Pawlowski, 2019; Riemann et al., 2020). The largest fraction of DR neurons are non-serotonergic (Descarries et al., 1982), express the VGAT marker (Allers and Sharp, 2003) and project to various forebrain targets including the LH (Bang and Commons, 2012). These DR_VGAT_ neurons control food intake and their activity increases in response to fasting (Nectow et al., 2017), however, their role in sleep-wake behavior remains to be examined.

Using combinatorial optogenetics and electrophysiological recordings from identified neurons, we show that LH_GABA_ neurons selectively inhibit DR_GABA_ neurons resulting in disinhibition of a part of DR neurons ultimately leading to arousal. We found that DR_GABA_ neurons are highly active during wakefulness compared to NREM and REM sleep. Selective activation of their projection to the LH area leads to wakefulness, highlighting the importance of reciprocal hypothalamus-brainstem long range circuits in the control of brain states and arousal.

## MATERIALS AND METHODS

All experimental procedures were performed according to the European Communities Council Directives of 1986 (86/609/EEC) and 2003 (2003/65/CE) for animal research and were approved by the Ethics Committee of the University of Szeged and University of Bern. 80 adult (age > 2 months) male mice were used in this study.

### Experimental design

#### Viral targeting

For *in vitro* recordings AAV1-CAG-ChR2-venus-WPRE-SV40, AAV1-CAGS-flex-ChR2-tdTomato-WPRE-SV40 viruses, (50-150 nl, Penn Vector Core; titer: 5×10^12^ gc/ml) were injected bilaterally in the LH (AP: −1.4 mm, ML: ±1 mm, DV: −5.0−5.4 mm) of WT, GAD-GFP (Tamamaki et al., 2003) or Vgat-IRES-Cre (Vong et al., 2011) mice using a nanoinjector (Nanoliter 2000, WPI) connected to a glass pipette (~20 um diameter) at a rate of 1 nl/s.

For *in vivo* optogenetic activation of LH_VGAT_ projections to the DR, one group of Vgat-IRES-Cre mice (Vong et al., 2011) received AAV1-CAGGS-FLEX-CHR2-td tomato-SV40 or AAV2-EF1a-DIO-hChR2(H134R)-eYFP, while the second group received AAV2-EF1a-DIO-eYFP (control, all virus vectors were packaged at the Vector Core of the University of North Carolina at Chapel Hill, titres between 3·10^12^ to 5·10^12^ gc/ml). For the intersectional optogenetic activation of DR neuronal subpopulations, Vgat-IRES-Cre (Vong et al., 2011) x Pet1-Flpe (Jensen et al., 2008) double transgenic mice received AAVdj-EF1a-Foff/Con-hChR2(H134R)-eYFP (Pet1-OFF/Vgat-ON), AAVdj-EF1a-Fon/Con-hChR2(H134R)-eYFP (Pet1-ON/Vgat-ON), or AAVdj-EF1a-eYFP (eYFP controls). 150-600 nl of each virus was injected bilaterally into the LH or into the dorsal raphe nucleus (AP: −4.45, ML: +1.2, DV: - 2.9 to −3.1, at a 30° angle). The virus suspension was infused with a micro-infusion pump (PHD Ultra, Harvard Apparatus) through a 28 G stainless steel cannula (Plastic One) at a rate of 0.1 μl per minute.

#### Surgical procedures

Mice were anesthetized using 1-1.5% (v/v) Isoflurane, placed in a stereotaxic frame (Model 900, David Kopf Instruments, USA) on a heating pad (Supertech, Hungary). For sleep studies 8-10 week old male Vgat-IRES-Cre mice were chronically implanted with an optical fiber (200 um, 0.39 NA Core Multimode Optical Fiber, FT200EMT, TECS Clad, Thorlabs, inserted into zirconia ferrules, 1,25 mm OD; 230 μm ID, Precision Fiber product) above the DR (AP −4.45, ML +1.2, DV −2.9 from Bregma at a 20° mediolateral angle), which were fixated to the skull with C&B-Metabond (Parkell Inc.). Four stainless steel screw electrodes were inserted epidurally (two into both frontal cortices, two into parietal cortices) for the recording of EEG signals and wire loops were sewed to both neck muscles for EMG recordings. The EEG/EMG electrodes and optic fiber were secured to the skull with dental cement (Patterson dental). After surgical procedures mice received appropriate post-operative care (analgesics: Rimadyl (15 mg/kg), anti-inflammatory drug: Dexafort (2mg/kg), antibiotic: Gentamicin (5mg/kg) and were allowed to recover for two weeks.

#### In vitro electrophysiology

Mice were deeply anesthetized and perfused through the heart with ice cold cutting solution containing (in mM) 93 *N*-methyl-d-glucamine, 2.5 KCl, 1.2 NaH_2_PO_4_, 30 NaHCO_3_, 20 HEPES, 25 glucose, 5 *N*-acetyl-l-cysteine, 5 Na-ascorbate, 3 Na-pyruvate, 10 MgSO4, and 0.5 CaCl_2_. The same solution was used to cut coronal brainstem slices containing the DR at 4°C and for the initial storage of slices (32-34°C for 12 minutes) following which the slices were stored in a solution containing (in mM) 30 NaCl, 2.5 KCl, 1.2 NaH_2_PO_4_, 1.3 NaHCO_3_, 20 HEPES, 25 glucose, 5 *N*-acetyl-l-cysteine, 5 Na-ascorbate, 3 Na-pyruvate, 3 CaCl2, and 1.5 MgSO_4_. For recording, slices were submerged in a chamber perfused with a warmed (34°C) continuously oxygenated (95% O_2_, 5% CO_2_) ACSF containing (in mM) 130 NaCl, 3.5 KCl, 1 KH_2_PO_4_, 24 NaHCO_3_, 1.5 MgSO_4_, 3 CaCl_2_, and 10 glucose. DR neurons were visualized using standard DIC optics and recorded in whole cell current clamp mode using an EPC9 amplifier (Heka Elektronik). Patch pipettes (tip resistance, 4–5 MΩ) were filled with an internal solution containing the following (in mM): 126 K-gluconate, 4 KCl, 4 ATP-Mg, 0.3 GTP-Na_2_, 10 HEPES, 10 kreatin-phosphate, and 8 biocytin, pH 7.25; osmolarity, 275 mOsm. The liquid junction potential (−13 mV) was corrected off-line. Access and series resistances were constantly monitored, and data from neurons with a >20% change from the initial value were discarded. DR_GABA_ neuron somata in GAD-GFP mice (Tamamaki et al., 2003) were targeted under epifluorescent illumination. Photostimulation was performed through the microscope objective using a blue LED light source (0.5-0.8 mW/mm^2^, Thorlabs).

#### In vivo electrophysiology and juxtacellular labeling

To record the activity of DR neurons mice were anesthetized using 1-1.5% (v/v) Isoflurane (Forane), placed in a stereotaxic frame (Model 900, David Kopf Instruments, USA) on a heating pad (Supertech, Hungary). A small craniotomy (1 mm diameter) was made over the DR (AP: −4.75, ML: 0.2, DV: −3.1 (mm) from Bregma) leaving the dura mater intact. Glass electrodes (resistance: 10-25 MOhm) were filled with saline containing 1.5% Biocytin (Sigma, Hungary). The electrode was lowered in a 15° mediolateral angle to avoid the sagittal sinus. The activity of DR neurons was monitored using a Multiclamp 700B amplifier (Molecular Devices) operating in current clamp mode (filtered between 100 Hz and 3 kHz). At the end of some recordings neurons were filled with Biocytin (Sigma Aldrich) using 0.5-4 nA anodal current pulses of 500 ms duration, 50% duty cycle for 2-10 minutes) as previously described (Pinault, 1996). For recording GABAergic neurons in the DR, the lateral wings were targeted, where most neurons are GABAergic (Allers and Sharp, 2003). These neurons possessed small (≤15 μm diameter) oval somata and mean baseline firing rates above 5 Hz (Allers and Sharp, 2003).

#### Data acquisition

Before the start of the experiment, the animals were habituated to a 3-m-long fiber optic patch cord with protective tubing (Thorlabs) that was connected to the chronically implanted optical fiber with a zirconia sleeve (Precision Fiber Products) and an EEG/EMG cable for one week. Subsequently, a 24-h baseline polysomnographic recording was obtained without any optogenetic manipulations. EEG and EMG signals were sampled with an AM systems 3500 amplifier at 512 Hz and digitized with a national instruments USB X DAQ device. EEG and EMG data were recorded with the SleepScore software (View Point). All experiments took place between 10 - 2 PM (Zeitgeber time 3-7).

For unit recordings in freely moving animals, mice were habituated to optical patch cords and an electrode board head stage (RHD2132, Intan Technologies) for at least 4 days. Subsequently a 24-h baseline polysomnographic recording was obtained without optogenetic manipulations to ensure that the animals recovered a normal sleep-wake cycle. Tetrode signals were acquired at 20 kHz with an open-source acquisition software (Open Ephys) via a digitizing head stage (RHD2132) and a multi-channel DAQ board (Open Ephys, Intan Technologies). Opto-tagging was performed by applying short blue light pulses to the DR (10 times 5ms pulses, every 30 seconds). Light-responsive cells were identified based on their spiking activity within 3 ms of the application of blue light.

#### EEG and EMG recordings

Sleep-wake stages were manually scored based on EEG/EMG signal characteristics as described previously (Jego et al., 2013). Briefly, wakefulness was scored when low-amplitude desynchronized EEG activity was accompanied by high, tonic muscle activity with phasic bursts in the EMG. NREM sleep was defined by the presence of synchronized, high amplitude oscillations in the delta frequency band (0.5 – 4 Hz) and low EMG tone. REM sleep was scored when the EEG showed pronounced theta oscillations with an almost complete absence of EMG tone except for short twitches. The 24 h baseline recordings were scored in epochs of 5 s, while state-specific optogenetic stimulation recordings were scored at an epoch duration of 1 s.

#### Optogenetic activation during polysomnographic recordings

For *in vivo* optogenetic stimulation experiments, the patch cord was connected to a 473nm DPSS laser (Laserglow Technologies) via a FC/PC connector (250 um, 30126G2-250, Thorlabs). The laser output was set to a power output of 10 mW from the tip of the optic fiber. Trains of 5 ms pulses at 5 or 20Hz were applied to LH_VGAT_ fibers or DR Pet1-OFF/Vgat-ON neurons and fibers using a Master-9 pulse generator (AM Systems). State-specific optical activation was performed based on visual inspection of online EEG and EMG signal parameters. For NREM and REM sleep-specific optogenetic activation, pulse trains were applied through the patch cord and optic fiber when the animal spent at least 10 seconds in NREM or REM sleep. The optogenetic stimulation was performed throughout the NREM or REM episode until the mouse transitioned to another state. Afterwards, the animal was left undisturbed to cycle between NREM sleep, REM sleep and wakefulness for 2-3 cycles before the next NREM or REM sleep episode was optically stimulated. A total of 9 NREM and REM sleep episodes per animal were optogenetically stimulated for each frequency (5 and 20 Hz).

#### Immunohistochemistry following sleep studies

At the end of the experiments, the mice were deeply anesthetized with an intraperitoneal injection of pentobarbital (250 mg/kg, Streuli Pharma) and transcardially perfused with 5 ml cold saline (0.9 % NaCl) followed by 20 ml 4% formaldehyde (Grogg Chemie). Subsequently, the brains were removed and stored in formaldehyde over night at 4 °C for post-fixation. The next day, the brains were transferred into phosphate-buffered saline (PBS) containing 30 % sucrose before freezing. The brains were cut into 30 μm sections with a cryostat (Hyrax C 25, Zeiss) and collected in PBS with 0.1 % Triton A-100 (Sigma-Aldrich), PBST.

For immunostaining, the sections were first incubated for 1 h at room temperature in a blocking solution containing 4 % bovine serum albumin (Sigma-Aldrich) in PBS-T. Then the tissue was incubated with anti-GFP (1:5000, A0262, Life technologies) and anti-Tph2 (1:400, ABN60, Merck) for 24 h. Afterwards, the sections were washed for 5 x 5 minutes in PBS-T and then incubated with DyLight 488 and AlexaFluor 555 secondary antibody (1:500 dilution, A96947 for eYFP, Life technologies; 1:500, A21428 for Tph2; Life technologies) at room temperature for 1 h. The sections were washed 2 x 10 minutes in PBS-T and 2-10 minutes in PBS and then mounted on glass slides. Cover slips were fixed on the slides with Fluoromount (F4680, Sigma-Aldrich). Photomicrographs were taken using a confocal microscope. For display, the image brightness and contrast was moderately adjusted in Adobe Photoshop CC.

#### Immunohistochemistry following in vitro recordings

Following *in vivo* electrophysiological recordings, slices were immersed in 4% paraformaldehyde (PFA) in 0.1□M phosphate buffer (PB; pH□=□7.4) at 4□°C for at least 12□h. After several washes with 0.1□M PB, slices were embedded in 10% gelatin, and further sectioned into slices of 50-70 μm thickness in the cold PB using a vibratome VT1000S (Leica Microsystems). Coronal sections were subjected to freeze-thaw over liquid nitrogen after cryoprotecting in 10% and 20% sucrose. The recorded cells were first visualized with incubation in Cy3-conjugated streptavidin (Jackson ImmunoResearch) for 2□h, diluted at 1:500 in tris buffer saline (TBS, SigmaAldrich). After examination by epifluorescence microscopy (Olympus BX60), the sections containing the soma of the labeled neurons were incubated in 10% normal horse serumin (NHS) in TBS to block non-specific antibody-binding sites. Free-floating sections were incubated in primary antibodies dissolved in TBS containing 0,1% Trinon X-100 ((TBS-Tx); Sigma Aldrich) for overnight at room temperature ((RT); 22□°C). The following primary antibodies were used: anti-5HT-Rb (1:1000, Jackson ImmunoResearch); anti-GABA-Rb (1:1000, Jackson ImmunoResearch). After several washes in TBS, the immunoreactions were visualized with Dylight649-DARb (1:400, Jackson ImmunoResearch) secondary antibodies. Finally, the sections were mounted in Vectashield mounting medium (VectorLaboratories). Images were taken by confocal microscope (Olympus FV1000).

#### Immunohistochemistry following in vivo single unit recordings

After experiments the animal was quickly perfused through the heart with 50 ml of PBS followed by 50 ml 4% PFA in 0.1 M PB. Following perfusion, brains were postfixed overnight and stored in 0.1 M PB with 0.05% sodium azide at 4°C as a preservative. The tissue was sectioned in PB at 50–70 μm on a vibratome (Leica VT 1000S). Coronal sections were permeabilized in 0.3 % TBS-Tx and incubated in Cy3 or Alexa488 conjugated streptavidin (1:500; Jackson ImmunoResearch) for 2h at RT in TBS. Images were taken by confocal microscope (Olympus FV1000).

### Statistical analysis

Data analysis was performed using Spike2 (Cambridge Electronic Design, UK), OriginPro 8.5 (MicroCal, USA), IgorPro (WaveMetrics, USA), Matlab (MathWorks, USA) and Prism (GraphPad, USA) software. The data were tested for normality with the Shapiro-Wilk and the Kolmogorov-Smirnov tests and accordingly analysed using parametric or non-parametric methods. We used three-way ANOVA to test for frequency (5 versus 20 Hz, within-group comparisons) and opsin (eYFP controls versus ChR2-expressing mice, between-groups comparisons) effects in the latencies to transition to another state from NREM or REM sleep of state-specific optogenetic activation experiments. PSPs evoked by photostimulation were detected in Vm recordings using a threshold of 2xSD in a ± 50 ms time window following photostimulation. Action potentials in extracellular recordings were detected in Spike2 using a straight forward level threshold. Firing rate histograms were computed for each photostimulation trial of each neuron. The control period was defined as 0.5 seconds before photostimulation. All data are displayed as averages +/- SEM.

The firing rate change of single units upon photostimulation was assessed by subtracting the average spike rate during the 10 seconds preceding the stimulation from the average spike rate during optogenetic activation. To test for significant modulation, 9999 stimulation onsets were randomly drawn from each single unit’s activity time series and the firing rate change across these random onsets was calculated. If the actual firing rate difference was larger than 95% or smaller than 5% of the randomly calculated ones, the firing of the neuron was significantly modulated during this photostimulation. Inhibited or disinhibited neurons showed a significant decrease or increase, respectively, of their firing rate during more than 50% of all photostimulations. Neurons whose firing rate was significantly modulated during less than 50% of all optogenetic stimulations are reported as unresponsive.

## RESULTS

### LH_GABA_ projections to the DR promote wakefulness

A subpopulation of LH_VGAT_ (vesicular GABA and glycine transporter) neurons causally promotes arousal from NREM sleep (Herrera et al., 2016; Venner et al., 2016; Venner et al., 2019). Here, we tested the role of LH_VGAT_ projections to the DR in the control of sleep-wake behaviour. To deliver opsins to LH_VGAT_ neurons, VGAT-Cre mice were stereotactically injected with AAV2-EF1a-DIO-eYFP (control) or AAV2-EF1a-DIO-hChR2(H134R)-eYFP) (ChR2-EYFP) in the LH for *in vivo* optogenetic activation experiments (see Materials and Methods; Figure 1A and B). Mice were then implanted with EEG/EMG electrodes for recording and quantification of sleep-wake states together with an optic fiber above the DR (see Materials and Methods; Figure 1C and D). State-specific optogenetic activation of LH_VGAT_ fibers in the DR was triggered manually after at least 10 seconds in stable NREM or REM sleep. Blue light (473 nm) was delivered to the DR at 5Hz or 20Hz (5ms pulses, see (Herrera et al., 2016)) in both control (eYFP) and ChR2-expressing animals continuously until a state transition occurred (Figure 1E). Interestingly, optogenetic activation of LH_VGAT_ projections to the DR during NREM sleep led to a rapid arousal at both 5 and 20 Hz within < 7s (Figure 1F, P < 0.0001, three-way ANOVA), whereas optical activation of these fibers failed to significantly change REM sleep duration (Figure 1G). These results extend previous mechanisms for LH_VGAT_ induced arousal to DR neurons in the brainstem.

**Figure 1.**
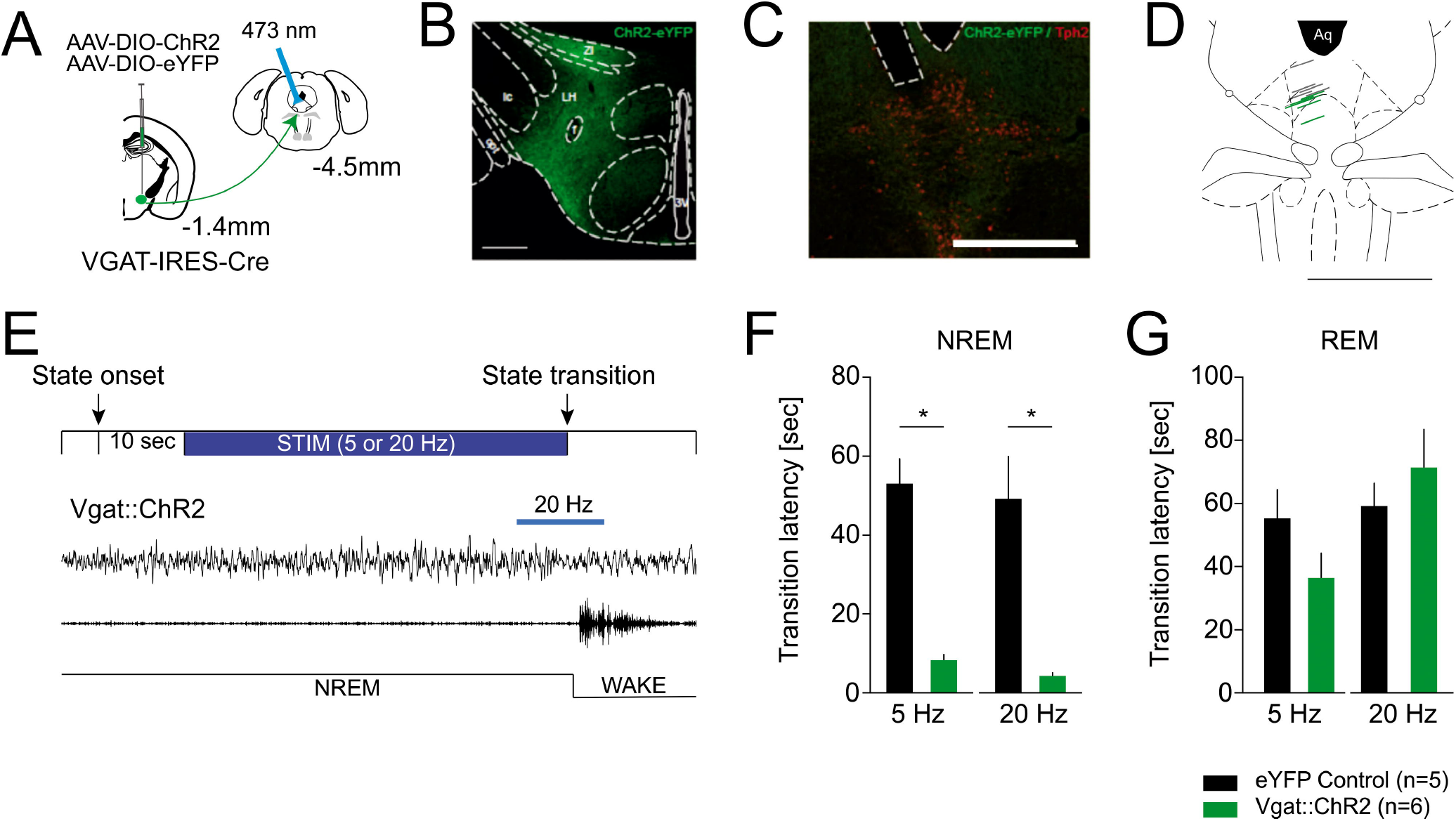
Stimulation of LH_GABA_ fibers in the DR promotes wakefulness. (A) Schematics of the experimental design: AAV-EF1a-DIO-ChR2-eYFP or AAV-EF1a-DIO-eYFP was injected in the LH of Vgat-cre mice. Following 4 weeks of expression LH_GABA_ fibers were photostimulated in the DR with simultaneous polysomnographic recordings for sleep stage scoring. (B) Example ChR2-eYFP expression in the LH. Fluorescent image overlaid on the stereotaxic atlas (AP: 4.45). (C) Location of the optical fiber in the DR used for photostimulation. Merged fluorescent image showing LH_GABA_ fibers expressing ChR2-eYFP (green) and TPH immunhistochemistry (red) to indicate the location of TPH synthetizing serotonergic neurons within the DR. (D) location of all implanted optical fibers in 11 mice. Green lines: Vgat::ChR2-expressing mice (n=6), gray lines: eYFP controls (n=5), Aq: aqueduct). (E) (Top) Schematic experimental timeline for sleep-stage specific photostimulation. (Bottom) Representative EEG/EMG traces illustrate behavioural response to optogenetic stimulation. Note the rapid EEG desynchronization and EMG activation upon photostimulation indicating a NREM-to-Wake transition. (F) Latency of NREM to wake transition caused by the photostimulation of LH_GABA_ axons in the DR. (G) Latency of REM to wake transition caused by the photostimulation of LH_GABA_ axons in the DR. * *P* < 0.05.

### DR_GABA_ neurons receive monosynaptic inputs from LH_GABA_ neurons

We previously found that LH_GABA_ neurons send long-range projections to various brainstem nuclei, including the DR (Herrera et al., 2016). In order to determine the downstream circuitries mediating this rapid arousal response, we mapped LH_GABA_ efferents utilizing ChR2-assisted circuit mapping to functionally characterize this LH-DR circuit (Figure 2A). We infected LH neurons irrespective of their neurochemical identity with ChR2 using a viral construct (AAV1-CAG-ChR2-venus-WPRE-SV40) optimised for in vitro photostimulation of axons (Petreanu et al., 2009). We found that photostimulation of ChR2 expressing LH terminals in the DR evoked inhibitory postsynaptic potentials (IPSPs, −7.9±1.03 mV peak amplitude with a 10,91±2,41 ms decay time) in 7 DR cells (14%) recorded (Figure 2DE), while EPSPs of 7.58±0.9 mV peak amplitude with a 11,07±2,46 ms decay time were recorded in 10 (20%) DR cells (Figure 2E). No detectable PSPs could be recorded in the remaining population (n=34, 66%). All DR neurons exhibiting IPSPs following LH axonal photostimulation were confirmed to be GABAergic (Figure 2B and E) by post-hoc immunohistochemistry (n=5) or by recording DR_GABA_ neurons in GAD-GFP mice (n=3, see Materials and Methods), one identified DR_GABA_ neuron received EPSPs (Figure 2E). The IPSPs could be abolished by SR95531 (i.e. gabazine, n= 5 out of 5 cells, 100%) but not NBQX (n=0/5, 0%, Figure 2D), suggesting that these are monosynaptic pathways implicating GABAa receptors. Note that this result was further confirmed by the recording of similar IPSPs upon photostimulation of LH axon terminals in the presence of TTX and 4-AP (mean IPSP amplitude: control: 6.5±0.6 mV, TTX/4-AP: 6.8±0.7 mV, n=6 neurons, Figure 2F and G). When neurons were held at membrane potentials close to action potential initiation threshold, photostimulation of LH axons also led to a prominent suppression of firing (12.66±6.65 Hz vs. 0 Hz, MI −1.0, n=3 neurons) due to the presence of IPSPs (Figure 2D). This suppression of firing was fully abolished when IPSPs were blocked by gabazine (n=3/3, Figure 2D). These results show that the LH to DR monosynaptic projections are functional, can be either inhibitory or excitatory, and that DR_GABA_ neurons are preferentially inhibited by LH_GABA_ fibers.

**Figure 2.**
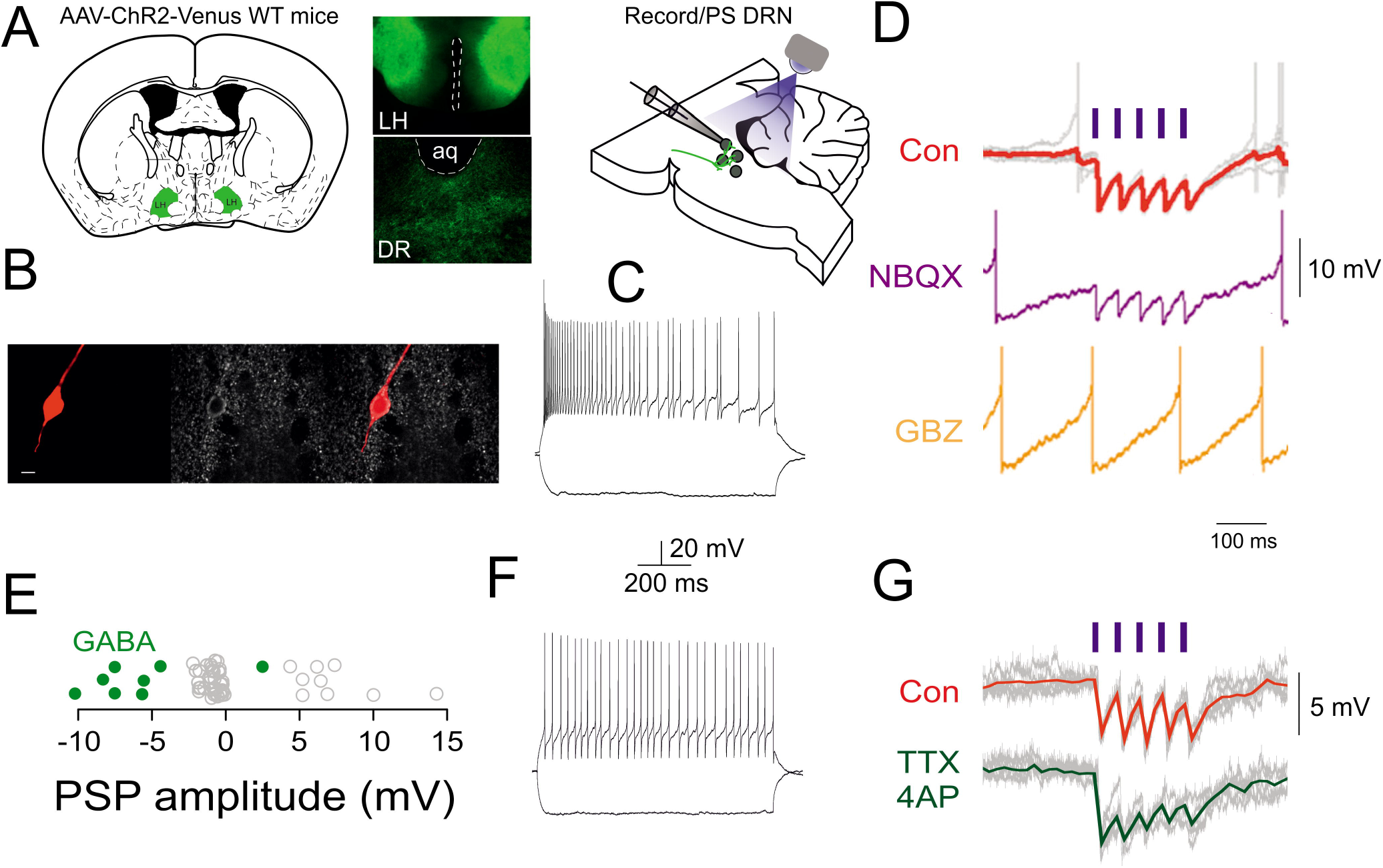
LH projections synaptically target DR neurons. (A) Schematic of the experimental design: AAV-ChR2-Venus injection in the LH of WT mice leads to prominent ChR_2_ expression in DR projecting axons. Four weeks after the infection local axons of LH neurons are photostimulated in the DR with simultaneous whole-cell patch clamp recordings in identified DR neurons. (B) Fluorescent images (Streptavidin, GABA immunostaining, merged) of a DR_GABA_ neuron responding to LH axonal photostimulation. (C) Membrane potential responses of the neuron in (B) to hyperpolarizing and depolarizing current steps shows fast action potential output with moderate frequency adaptation. (D) The photostimulation of LH axons leads to monosynaptic IPSPs that are resistant to iGluR blockade (NBQX), but blocked by SR95531 (i.e. gabazine, GBZ). (E) Distribution of PSP amplitudes in all the neurons recorded (n=51), green circles indicate identified DR_GABA_ neurons. (F) Membrane potential responses of a physiologically identified DR_GABA_ neuron. (G) The photostimulation of LH axons leads to IPSPs in the neuron shown in (F) that persist following the application of TTX and 4-AP.

### LH_GABA_ to DR projections can rapidly suppress the activity of DR_GABA_ neurons

To test for functional connections from LH_GABA_ projections to DR neurons, we recorded extracellular single unit activity from morphologically identified neurons in the DR of anesthetized VGAT-cre mice infected with AAV1-CAGGS-FLEX-CHR2-tdTOM.-SV40 in the LH while photo-stimulating ChR2 expressing LH_GABA_ axons in the DR (5 light pulses of 10 ms at 20 Hz, 5 mW; Figure 3A).

**Figure 3.**
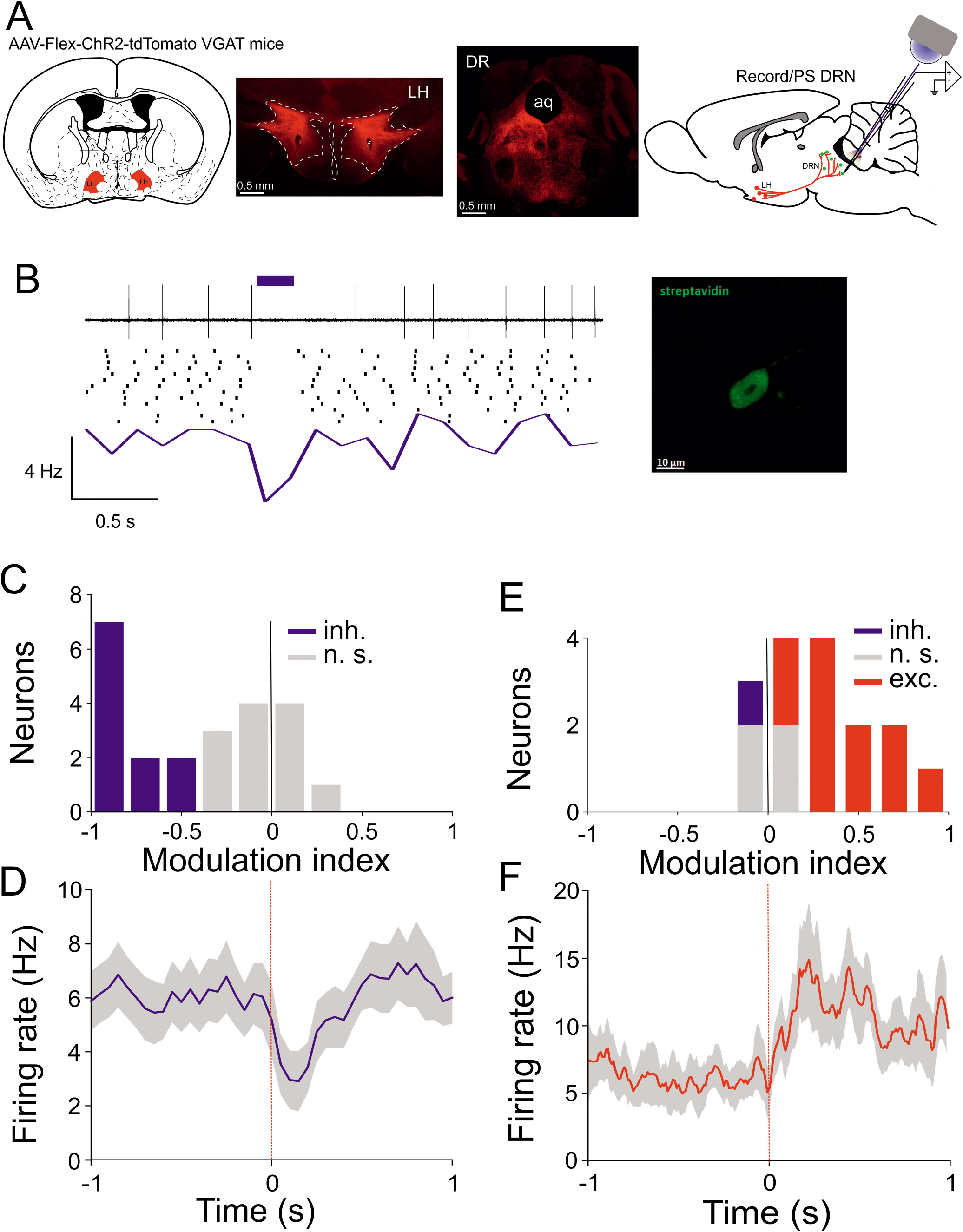
LH_GABA_ inputs control DR neuronal activity. (A) Schematic of the experimental design: AAV-Flex-ChR2-td tomato injection in the LH of VGAT mice leads to prominent ChR2 expression in DR projecting axons. 4 weeks after the infection local axons of LH_GABA_ neurons are photostimulated in the DR with simultaneous single unit juxtacellular recoding/labeling. (B) Example of spontaneous activity of a DR_GABA_ neuron and its response to local photostimulation of LH_GABA_ axons. (Top) Single-trial raw data aligned on photostimulation onset (blue horizontal bar, 5 mW, 10 ms pulses at 50 Hz). Middle, Raster plot in which each tic is a single-unit spike and each row represents a single trial. Red line: photostimulation onset. (Bottom) PSTH of the unit shown above. (Right) Fluorescent pictures showing the neuron recorded in (B) filled with Neurobiotin using the juxtacellular filling technique. Left: streptavidin, middle: VGAT immunostaining, right: merged picture. (C) Modulation index of all recorded neurons under anaesthesia. (D) Grand average PSTH of all neurons recorded neurons under anaesthesia. (E) Modulation index of all recorded neurons in awake animals (inh: inhibited, n.s.: not significant, exc: excited). (F) Grand average PSTH of all neurons recorded neurons in awake animals.

When comparing the activity of DR neurons upon LH_GABA_ axonal photostimulation, we found a rapid suppression (peak suppression: 42.37±5.94%, Figure 3B) of neuronal activity (baseline firing: 7.72±6.43 Hz, baseline firing rate after photostimulation: 5.37±6.77 Hz, n=25, *p* < 0.001, Wilcoxon’s signed rank test, Figure 3C and D). We identified the morphology of 8 neurons recorded and filled with the juxtacellular labeling technique. These neurons were classified as putative DR_GABA_ neurons based on their high baseline firing rate (≥6 Hz) morphology and location within the DR (see Materials and Methods). These results show that LH_GABA_ neurons exert an inhibitory action onto putative DR_GABA_ neurons.

We next explored the effect of stimulating LH_GABA_ fibers in the DR in awake, head-restrained animals while recording the activity of multiple single units in the DR (see Methods). Comparison of the activity of DR neurons recorded in the presence and absence of LH_GABA_ axonal photostimulation (5 light flashes of 10 ms at 20 Hz, 5 mW) further confirmed the suppressive effect in a subset of DR neurons (2/12, 17%), while the activity of the remaining neurons was increased (10/12, 83%). The overall activity of DR neurons was slowly (~200 ms), but persistently (~1 sec) increased (baseline firing: 7.67±6.87 Hz, baseline firing after photostimulation: 10.98±8.920 Hz, n=12, *p* < 0.001, Wilcoxon’s signed rank test, Figure 3E and F).

### The activity of identified DR_GABA_ neurons is brain state dependent

To target DR_GABA_ neurons in the DR, we made use of the intersectional virus approach (Fenno et al., 2014). In brief, the INTRSECT system (‘intronic recombinase sites enabling combinatorial targeting’) consists of orthogonal Cre and Flp recognitions sites (lox and FRT sites, respectively) that are inserted into synthetic introns of the ChR2-eYFP open reading frame (ORF) within an adeno-associated viral construct (AAVdj). The starting direction of the ORF fragments between the lox and FRT recognition sites defines whether the presence of Flp and/or Cre recombinases is required to produce a functional opsin (ChR2-eYFP or eYFP, see Figure 4A and B).

**Figure 4.**
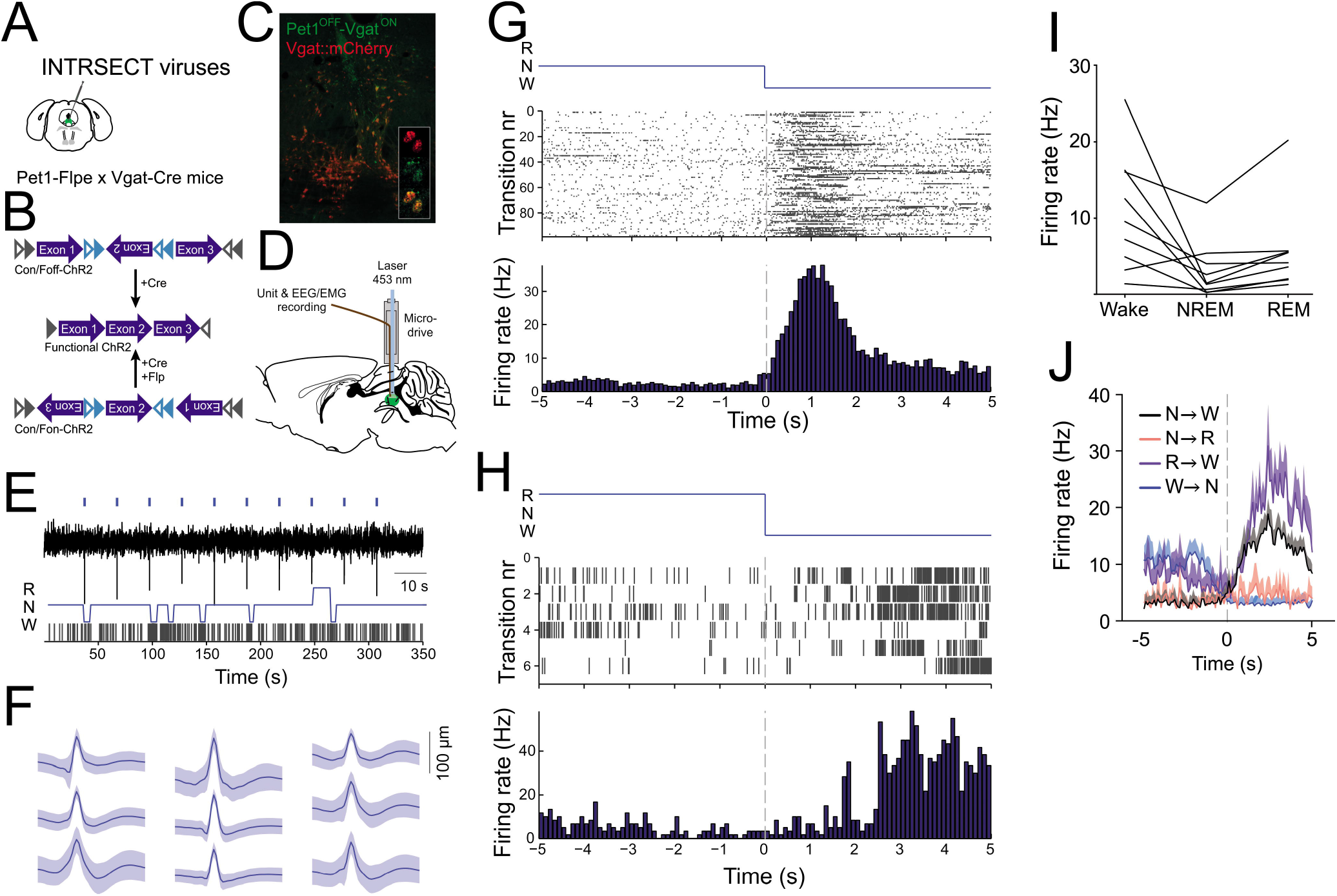
The activity of optogenetically identified DR_GABA_ neurons are modulated across the sleep-wake cycle. (A) Schematics of intersectional viral strategy: ChR2-Con/off-eYFP was targeted to DR neurons in Pet1^Flpe^-Vgat^Cre^ double transgenic mice. (B) Schematics of INTRSECT system (Fenno et al., 2014): FRT (grey triangles) and lox sites (light blue triangles) were introduced in intronic sequences within the ChR2 ORF allowing for directional control of these ORFs. In the Con/Foff-ChR2 construct, the presence of Cre recombinase produces a functional ChR2. However, the presence of Flp recombinase would invert the exons, thus yielding an inactivated state of ChR2. In the Con/Fon-ChR2 construct, the presence of both Cre and Flp recombinase is required to produce a functional ChR2. (C) ChR2 expression is restricted to DR_GABA_ (Vgat-mCherry expressing) neurons. (D) Schematics of the experimental design: simultaneous DR Pet1-OFF/Vgat-ON neuron and and EEG recordings in behaving mice. (E) Example recording of a DR Pet1-OFF/Vgat-ON neuron throughout various sleep/wake states. (Top) raw trace voltage recording shows clear action potential firing upon photostimulation (blue ticks). (Bottom) Hypnogram and spike ticks of the neuron showed above (W: wake, R: REM, N: NREM). (F) Averaged spike waveforms of optogenetically identified DR Pet1-OFF/Vgat-ON neurons (n = 9 neurons from n = 4 mice). DR Pet1-OFF/Vgat-ON neurons were identified by their immediate firing in response to pulses of blue light generated by a 453 nm laser (10 pulses of 5ms every 30 sec). (G, H) Raster plot of a DR Pet1-OFF/Vgat-ON neuron during NREM-to-wake transitions (G) and REM-to-wake transitions (H) with spike ticks of the neuron (Top) and PSTH of the neuron across all transitions (Bottom). (I) Firing rates of the 9 DR Pet1-OFF/Vgat-ON neurons across wakefulness, NREM and REM sleep. (J) PSTH (Mean + SEM) of DR Pet1-OFF/Vgat-ON neurons during the different sleep state transitions.

Here, we used this approach to exclude the possibility that GABAergic markers (Vgat) are co-expressed with serotonergic neurons (Huang et al., 2019; Ren et al., 2019; Okaty et al., 2020). To this end, Pet1^Flpe^ - Vgat^Cre^ double transgenic mice (Vong et al., 2011; Jensen et al., 2008) were injected with Cre-ON/Flp-OFF or Cre-ON/Flp-ON intersectional viruses (Figure 4A and B), which allowed the targeting of ChR2-eYFP to distinct subpopulations of the DR that expressed Pet1, a transcription factor predominantly found in 5-HT neurons (Hendricks et al., 1999), and Vgat (Pet1-ON/Vgat-ON) or Vgat only (Pet1-OFF/Vgat-ON). The Pet1-OFF/Vgat-ON GFP-positive neurons co-localized with Vgat neurons that were co-transfected with a Cre-dependent mCherry virus (Figure 4C).

To investigate whether the activity of DR_GABA_ neurons is modulated across sleep-wake states, we performed opto-tag unit recording of putative DR Pet1-OFF/Vgat-ON neurons while simultaneously conducting EEG and EMG recordings in freely moving Pet1^Flpe^-Vgat^Cre^ mice (see Materials and Methods; Figure 4D). These neurons were identified by their immediate increase in spiking in response to short blue light pulses (10 times 5ms pulses, every 30 seconds) delivered to the DR via an optic fiber. A total of 9 optogenetically identified Pet1-OFF/Vgat-ON neurons (9 out of 66 cells from 4 animals; Figure 4E and F) were recorded across the sleep-wake cycle (Figure 4F). The activity of Pet1-OFF/VGAT-ON neurons was modulated across sleep-wake states (Figure 4G-I, NREM vs Wake/REM: P < 0.05, Friedman non-parametric test). Indeed, their firing activity increased significantly upon arousal as compared to the 5 seconds before the transition occurred (i.e., when the muscle tone was still low; Figure 4G, H and J; NREM-to-wake P < 0.01; REM-to-wake P < 0.05, paired t-tests). In turn, the firing activity decreased when the mice were at the onset of NREM sleep (Wake-to-NREM P < 0.01, paired t-test), while on the other hand, the transition from NREM to REM sleep was not marked by a substantial change in firing activity (Figure 4J).

### Wake promoting effect of DR_GABA_ neurons

Next, we investigated whether Pet1-OFF/VGAT-ON neurons in the DR were causally involved in arousal control. For this, we optogenetically activated DR Pet1-OFF/Vgat-ON neurons during NREM sleep (see methods; Figure 5C). Consistent with our correlative data, we found that photostimulation of DR Pet1-OFF/VGAT-ON neurons induced a rapid arousal from NREM sleep when light pulses were delivered at 20Hz as compared to controls (P < 0.0001, Three-way ANOVA; Pet1-OFF/VGAT-ON vs Ctrl: P < 0.0001, Dunnett’s multiple comparison’s post-hoc test), whereas 5Hz stimulation frequency did not decrease the latency to wakefulness transitions (n = 6 animals per group; Figure 5C). The probability of REM sleep onset decreased upon optogenetic activation of Pet1-OFF/VGAT-ON neurons during NREM sleep (Figure 5E), suggesting an implication in REM sleep onset. However, both 5 and 20Hz stimulation frequencies during REM sleep led to an immediate arousal from REM sleep in Pet1-OFF/Vgat-ON mice (Figure 5D, P < 0.0001, three-way ANOVA; Pet1-OFF/VGAT-ON vs Ctrl: P < 0.0001, at both 5 and 20 Hz, Dunnett’s multiple comparison’s post-hoc test). As expected, optogenetic activation of Pet1-ON/VGAT-ON neurons (n = 6 animals) did not alter sleep-wake behavior as compared to controls (n = 6 animals; Figure 5D, E).

**Figure 5.**
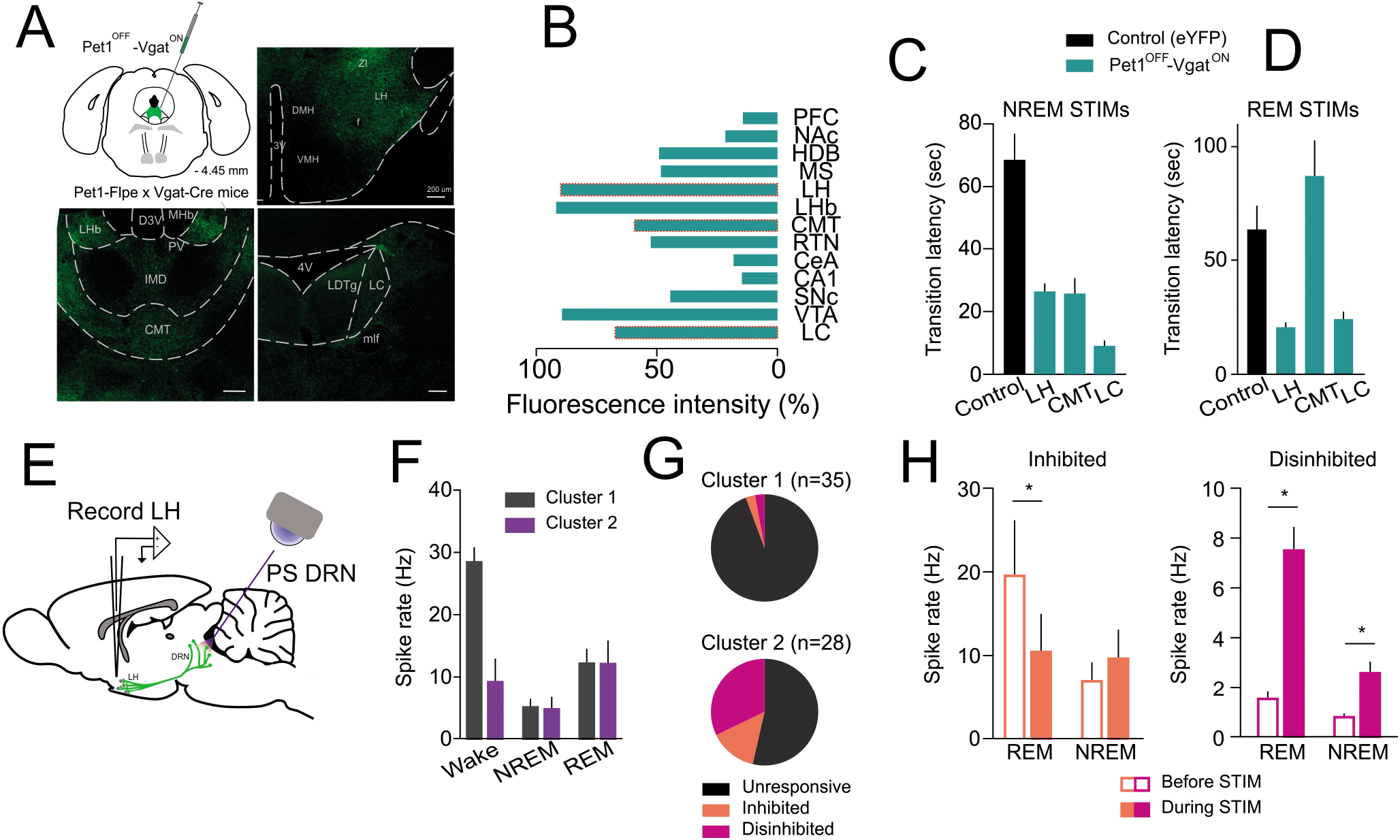
Stimulation of DR_GABA_ neurons promotes wakefulness. (A) Schematic of the experimental strategy: various intersectional viral constructs carrying the coding sequence of either eYFP (control) or ChR2 are injected in the DR of Pet1^Flpe^ - VGAT^Cre^ double transgenic mice. The DR is photostimulated 4 weeks post-infection. (B) Location of the fiberoptic canulae in all the animals used: left, gray lines: control, middle, green lines: Pet1-OFF/VGAT-ON, right, blue lines: Pet1-ON/VGAT-ON. (C) Latency of NREM to wake transition caused by the photostimulation of the DR. (D) Latency of REM to wake transition caused by the photostimulation of the DR. (E) State transition probabilities at 5 versus 20 Hz photostimulation for all three viral constructs. * *P* < 0.05.

### DR_GABA_ neurons promote wakefulness through disinhibition in LH

To identify potential sleep-wake centers that may mediate the arousal promotion of DR_GABA_ neurons, we performed a ChR2-assisted functional circuit mapping. Animals were prepared as described above. We found that DR Pet1-OFF/VGAT-ON strongly innervated major sleep-wake centers, such as the locus coeruleus (LC), the ventral tegmental area (VTA), the centromedial thalamic nucleus (CMT), and the LH (Figure 6A and B). To further understand the reciprocal connection between the DR and the LH, we performed optogenetic state-specific stimulations of Pet1-OFF/VGAT-ON projecting fibers in the LH. We found that optogenetic stimulation (20 Hz) of DR Pet1-OFF/VGAT-ON projections to the LH reproduced the arousal-inducing effect that was observed during somatic optogenetic activation (Figure 6C, D, NREM sleep: P < 0.0001, three-way ANOVA; Ctrl. versus LH stimulation: P < 0.001, Dunnett’s multiple comparisons post-hoc test; REM sleep: P < 0.0001, three-way ANOVA; Ctrl. versus LH stimulation: P < 0.001, Dunnett’s multiple comparisons post-hoc test). Interestingly, optogenetic activation of Pet1-OFF/VGAT-ON projections in the LC area, a predominant wake-promoting center, resulted in a similar effect (Ctrl. versus LC stimulation: NREM sleep: P< 0.0001; REM sleep: P < 0.001, Dunnett’s multiple comparisons post-hoc test), while optogenetic stimulation of Pet1-OFF/VGAT-ON projections in the CMT led to an arousal only from NREM sleep, but not from REM sleep (Ctrl. versus CMT stimulation: NREM sleep: P< 0.001; REM sleep: n.s., Dunnett’s multiple comparisons post-hoc test), in line with a previous study highlighting a role of the CMT in NREM sleep regulation (Gent et al., 2018) (Figure 6C, D).

**Figure 6.**
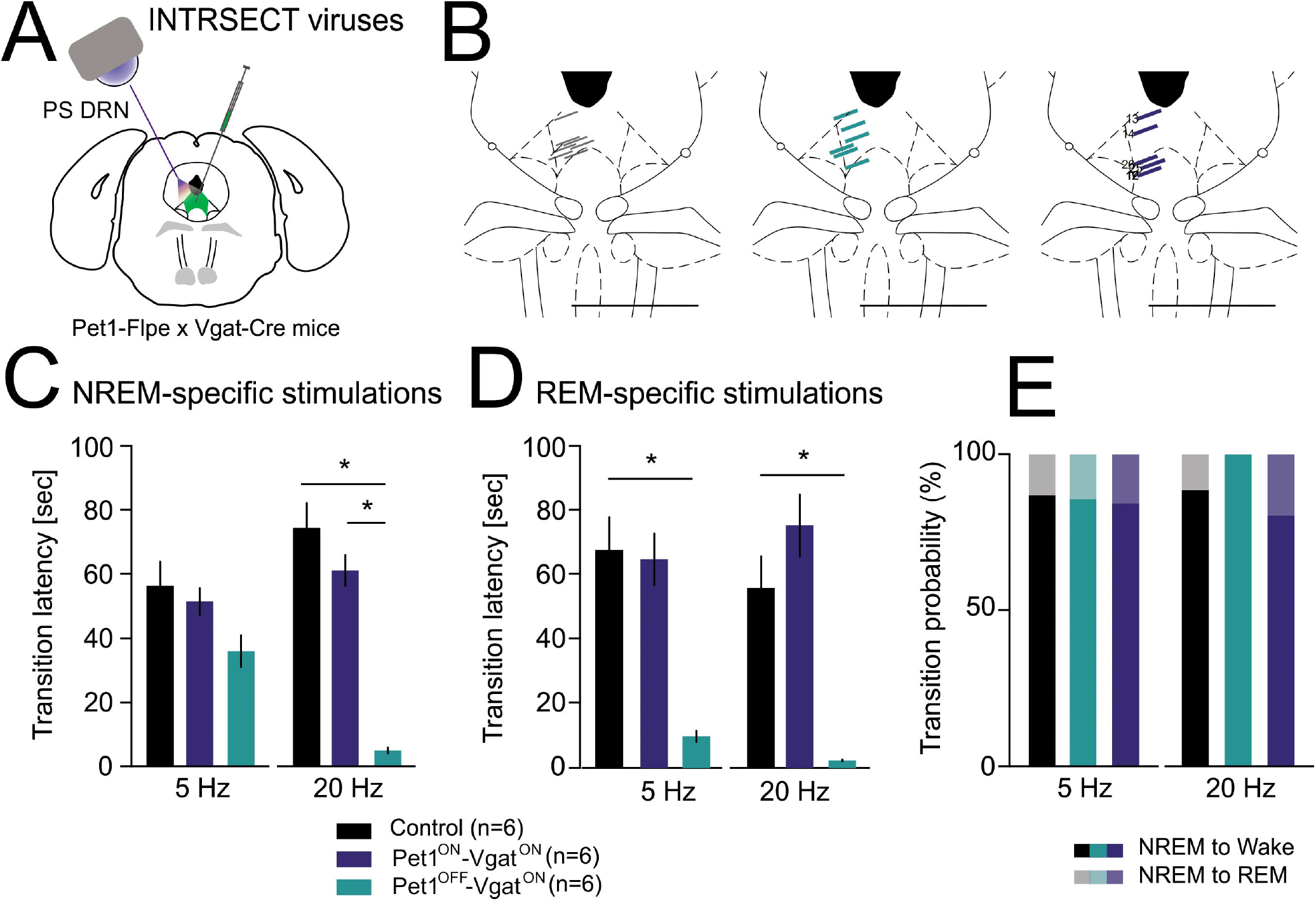
DR_GABA_ neurons promote arousal through disinhibition in the LH. (A) (Top left) Schematic of the experimental design: ChR2 is expressed in DR_GABA_ neurons following a viral injection in Pet1-Flpe x VGAT-Cre mice. Axonal projections of DR_GABA_ neurons express ChR2 in various DR target areas including the LH (top right), centromedial thalamus (CMT, bottom left) and locus coeruleus (LC, bottom right). (B) Normalized intensity of the fluorescent eYFP signal in various projection areas of the DR (PFC: prefrontal cortex, NAc: nucleus accumbens, HDB: horizontal diagonal band of Brocca, MS: medial septum, LHb: lateral habenula, RTN: thalamic reticular nucleus, CeA: central amygdala, CA1: hippocampus CA1 region, SNc: substantia nigra pars compacta). (C) Latency of NREM to wake transition caused by photostimulation of DR_GABA_ projections in the LH, CMT and LC. (D) Latency of REM to wake transition caused by photostimulation of DR_GABA_ projection targets (LH, CMT, LC). (E) Schematic of the experimental design: ChR2 expressing Pet1-OFF/Vgat-ON neurons are photostimulated in the DR while single units are recorded in the LH of freely moving mice. (F) Activity of all recorded LH neurons during various sleep-wake stages differentiates two clusters of neurons: Cluster 1 containing wake active neurons while Cluster 2 neurons do not show significant firing rate increases during the awake state. (G) Pie chart of all Cluster 1 and 2 LH neurons constructed on the basis of their response to the photostimulation of DR_GABA_ neurons. (H) Effect of DR_GABA_ photostimulation on the state-dependent firing of inhibited (left) and disinhibited (right) neurons. For details see main text. * *P* < 0.05.

To further delineate the mechanism by which DR Pet1-OFF/VGAT-ON neurons induce arousal via the LH, we performed single unit recordings in the LH while simultaneously recording EEG/EMG and conducting optical stimulations of ChR2-expressing Pet1-OFF/VGAT-ON neurons using a similar preparation as described above (see Materials and Methods; Figure 6E). First, we clustered all the cells that we recorded in the LH according to their sleep-wake discharge profiles; recorded cells separated into 2 clusters, with one cluster showing maximal activity during wakefulness (55% of cells from 2 animals; Figure 6F, G), while the second cluster showed maximal activity during both REM sleep and wakefulness (45% of cells from 2 animals; Figure 6F, G). Cells from both clusters were less active during NREM sleep (Figure 6F). Interestingly, during optogenetic stimulation of DR Pet1-OFF/VGAT-ON neurons, cells from Cluster 1 barely changed their firing rate, whereas half of the cells from the second cluster significantly modulated their firing activity in response to DR Pet1-OFF/VGAT-ON neuronal activation (Figure 6G). In particular, a major part of responsive cells from Cluster 2 was disinhibited by the optogenetic activation of DR Pet1-OFF/VGAT ON neurons, while a few cells were inhibited (Figure 5G and H, P < 0.05, three-way ANOVA). These results show that the activation of DR_GABA_ neurons disinhibits the majority of LH neurons and promotes wakefulness.

## Discussion

In this study, we show that activation of LH_GABA_ axons in the DR promote wakefulness through a local release of GABA that rapidly suppresses activity of identified DR_GABA_ neurons but indirectly activates other non-GABAergic DR neurons. In addition, we found that the activity of DR_GABA_ neurons is state dependent, while activation of DR_GABA_ neurons disinhibits the majority of LH neurons to promote wakefulness. This wake promoting effect of optogenetic activation of the LH_GABA_-DR_GABA_ circuit promotes rapid wakefulness selectively during NREM sleep, consistent with their activity profile across sleep-wake states (Hassani et al., 2010; Herrera et al., 2016). These findings identify a LH_GABA_-DR_GABA_ feedback loop that is essential for rapid alteration of global brain states and facilitation of wakefulness.

Using viral tracing and *in vitro* ChR_2_ assisted circuit mapping we show that a subset of LH_GABA_ neurons establishes monosynaptic connections to DR neurons. Functional analysis of this mapping revealed that LH_GABA_ neurons exert a GABAA-mediated inhibitory action on DR_GABA_ neurons leading to rapid NREM sleep-to-Wake transitions. The precise neurochemical identity of this subset of DR-projecting LH_GABA_ neurons remains to be established in light of the numerous sub-populations amongst inhibitory neurons of the LH (Mickelsen et al., 2017).

Local GABA release evoked by optogenetic activation of LH_GABA_ axons in the DR promotes wakefulness suppressing the activity of DR_GABA_ neurons. This occurs through a direct synaptic inhibition of DR_GABA_ neurons mediated by GABA_A_ receptors and leads to a prominent suppression of firing in DR_GABA_ neurons *in vivo*. Consistent with these *in vivo* recordings we show that the activity of identified DR_GABA_ cells is strongly modulated across NREM sleep-to-wake transitions and optogenetic activation of the LH_GABA_-DR circuit promotes rapid wakefulness selectively during NREM sleep, but not during REM sleep. This result is in line with previous studies reporting sleep/wake transitions upon optogenetic activation of LH_GABA_ fibers in the medial septum, locus coeruleus, reticular thalamic nucleus and ventrolateral preoptic area (Herrera et al., 2016; Venner et al., 2016; Venner et al., 2019). Our results demonstrate that LH_GABA_ neurons directly inhibit DR_GABA_ neurons and this probably results in a disinhibition of various DR neurons. Revealing the exact targets of this disinhibition awaits further investigation and are hard to predict given the extreme neurochemical and physiological diversity of the raphe nuclei (Domonkos et al., 2016; Szonyi et al., 2016; Sos et al., 2017). A phasic increase in 5-HT neuron activity can rapidly and prominently affect cortical activity (Lottem et al., 2016) and thus may lead to brain state changes. However, recently the selective tonic activation of DR_5-HT_ neurons has been shown to promote sleep, while burst stimulation induces wakefulness (Oikonomou et al., 2019). In light of the present results, it is possible that the phasic activity changes of DR neurons and their consequences at the cortical level at least partly originate from the LH_GABA_ neuron activation during sleep to wake transitions.

DR_GABA_ neuronal stimulation also promoted wakefulness, possibly by suppressing or activating local sleep- or wake-promoting neurons in the LH, respectively. One may speculate that GABA release from DR_GABA_ synapses could suppress the activity of sleep-promoting neurons including REM-sleep promoting MCH neurons (Jego et al., 2013; Konadhode et al., 2013; Tsunematsu et al., 2014), REM active LH_VGAT_ neurons (Hassani et al., 2010; Oesch et al., 2020), REM-active Lh6x GABA neurons from the ventral zona incerta (Liu et al., 2017). In contrast, it could inhibit local inhibitory circuits that would result in the activation of the wake-promoting hypocretins/orexins neurons (Adamantidis et al., 2007), or wake-active LH_VGAT_ neurons (Hassani et al., 2010; Herrera et al., 2016; Oesch et al., 2020). Whether and how this dual modulatory activity operates during the sleep-wake cycle awaits further investigation.

Taken together, a brain area involved in energy balance and arousal can affect the activity of the DR, an area involved in the control of higher brain functions including reward, patience, mood and sensory coding suggesting a potential route for interactions between metabolic states and brain states.

## Acknowledgements

This work has been supported by the Hungarian Scientific Research Fund (grant NF 123831 to MLL), the Hungarian Brain Research Program (grant KTIA_NAP_13-2-2014-0014 to MLL), NIH R01 DA034022 (SMD) and the German Research Foundation (GA 2410/1-1 to MG), the Human Frontier Science Program RGY0076/2012 (AA), Swiss National Science Foundation (grant 31003A_156156 to AA), European Research Council (grant 725850 to AA). MLL is a grantee of the János Bolyai Fellowship.

## Notes

### Competing Interest Statement

The authors have declared no competing interest.

